# Patterns of intersubject correlations parallel organizational gradients during naturalistic viewing

**DOI:** 10.1101/2025.10.19.683222

**Authors:** Meaghan Smith, Ahmad Samara, Donna Gift Cabalo, Alexander Ngo, Hallee Shearer, Tamara Vanderwal, Boris Bernhardt

## Abstract

Recent studies have robustly demonstrated that human cortical function can be described through sensory-transmodal gradients of cortical function, while naturalistic movie watching paradigms have been leveraged to index cortical synchronization. We leveraged two independent 7T movie fMRI datasets to assess correlations between intersubject correlation and functional gradients across movies, datasets, and spatial scales. At the whole-brain level, we observed robust relationships between intersubject correlations and a visual-transmodal connectivity gradient which was independent of movie content. Within functional networks, correlations were particularly pronounced for the visual, dorsal attention, and default mode networks. Our results demonstrate that naturalistic paradigms can provide targeted insight into multiscale processing hierarchies. Robust relationships across movies suggest that movie-watching can be viewed as a brain state that is independent of movie-content. Overall, this work suggests an important confluence of within-subject functional organizational axes and inter-subject synchronization when the brain is engaged in the processing of naturalistic stimuli.

## 1. INTRODUCTION

Two foundational goals of neuroscience are to determine the organizational principles that govern brain function and to further understand what makes each brain unique. To this end, imaging modalities such as functional magnetic resonance imaging (fMRI), along with the vast array of techniques leveraged to interpret the associated blood-oxygen level dependent (BOLD) signal, are powerful tools for investigation. By exploring differences in fMRI signal both across brain regions within individual brains, and within regions across brains, various analytic approaches to fMRI data have helped build our understanding of the brain’s functional organization (Biswal et al., 1995; Hasson et al., 2004). Recent research in the field of fMRI has complemented traditional task-based and task-free “resting-state” paradigms with naturalistic paradigms where participants watch movies while in the scanner (Eickhoff et al., 2020; Finn & Bandettini, 2021; Meer et al., 2020; Vanderwal et al., 2019). By eliciting robust, whole-brain responses with regions of systematic synchronization across subjects (Hasson et al., 2004; Jääskeläinen et al., 2008; Kauppi et al., 2010), this method is well suited to investigate the relationship between intersubject and inter-region variability in the brain. This innovation presents a powerful tool to probe brain function under conditions that more closely resemble everyday life, while also allowing us to improve both data quantity and quality by decreasing head motion in the scanner (Eickhoff et al., 2020; Frew et al., 2022; Vanderwal et al., 2015). Movie-watching additionally serves to synchronize low-level sensory processing, making key individual differences more salient (Vanderwal et al., 2017). By driving the brain in a way that is both more targeted and less constrained, movie fMRI gives us access to key methodological advantages of both traditional task-based fMRI and resting-state fMRI (Finn & Bandettini, 2021; Nastase et al., 2020; Sonkusare et al., 2019; Vanderwal et al., 2019).

Functional connectivity (FC) is defined as the correlation between fMRI activity over time across every pair of regions in the brain (Biswal et al., 1995; Menon & Krishnamurthy, 2019). Unsupervised learning algorithms represent a powerful tool to extract latent patterns from FC data, which can be difficult to interpret when studied in their high dimensional form. Non-linear dimensionality reduction identifies spatial eigenvectors–also termed gradients–of connectivity variation across the brain (Coifman & Lafon, 2006; Ferguson et al., 2011; Margulies et al., 2016; Paquola, Wael, et al., 2019; van der Maaten et al., 2007). These tools recently revealed a principal gradient of cortical organization that describes an evolutionarily conserved neural hierarchy (Margulies et al., 2016; Xu et al., 2020, p. 202). At one end of this principal gradient, we find unimodal sensory processing systems, while the other end is anchored by transmodal regions that serve high-level functions such as integration of sensory input and more abstract representations of information (Bernhardt et al., 2022; Burt et al., 2018; Mesulam, 1998; Paquola, Wael, et al., 2019; Smallwood et al., 2021). Beyond the functional significance of these gradients, it was demonstrated that regions that were maximally separated along this principal gradient were also maximally spatially segregated within the brain (Margulies et al., 2016). While the foundational gradient mapping work has typically studied task-free functional MRI data, recent advances have used naturalistic stimuli to investigate large-scale functional organization while the brain is active and processing complex multimodal stimuli. Extracted from movie-watching fMRI, these gradients are highly reliable across different movies and may show better correlation with behavioural measures than traditional resting-state gradients (Samara et al., 2023).

Beyond the characterization of brain functional organization via fMRI signal response amplitude and FC, it is also informative to investigate response reliability across individuals. Intersubject correlation (ISC) is a specific instance of response reliability analysis which leverages the time-locked nature of movie stimuli and allows us to collapse fMRI signal across time and across individuals, yielding a meaningful group-level measure of similarity at each brain region. These correlations have been shown to be unexpectedly high in certain brain areas when computed in subjects scanned during movie-watching (Hasson et al., 2004; Jääskeläinen et al., 2008; Kauppi et al., 2010; Nastase et al., 2019). High ISCs are detected in auditory and visual cortices, as well as brain regions involved in recognition of faces, objects, and buildings. Follow-up studies presented evidence suggesting that those regions which did not produce reliable ISCs could be considered to make up an ‘intrinsic’ network which has significant overlap with the default-mode network (DMN) (Buckner et al., 2008; Golland et al., 2007; Paquola et al., 2025; Smallwood et al., 2021). These results suggest that ISCs may represent a processing gradient between sensory and higher order regions which parallels the main axis identified by gradient analyses.

Recent literature has also indicated that ISCs may be able to capture brain-behaviour relationships where other measures are not. For example, children whose whole-brain neural activity has higher correlations with average whole-brain activity in adults during naturalistic viewing had higher performance scores in mathematics assessments while conventional task-based approaches were nonsignificant (Cantlon & Li, 2013). It has additionally been demonstrated that ISCs are significantly correlated with the degree of similarity between depressive symptom profiles in adolescents viewing emotional naturalistic stimuli (Gruskin et al., 2020). These results indicate that ISC could be sensitive to normative functional neurodevelopment in ways that conventional analyses of the BOLD-signal are not, suggesting potential utility as a brain-based biomarker. It thus becomes even more pressing to improve our understanding of these measures, and how they might relate to well-established FC patterns.

The current study uses high field fMRI data acquired at 7T to detect relationships between whole-brain organizational principles that emerge within individuals, and stimulus-locked response reliability patterns that emerge across individuals. In using these two measures in parallel, we aim to better understand how intersubject and inter-regional variability co-occur in the brain. Additionally, leveraging two data-driven approaches allows us to identify latent patterns in the data without imposing any existing assumptions about brain dynamics. Here, we use movie fMRI data to compute gradient and ISC scores at the individual and group level to determine if a meaningful spatial relationship exists between these two measures. Using 7T movie fMRI data from the Human Connectome Project (HCP) (Van Essen et al., 2013) as well as the Precision Neuroimaging (PNI) fMRI data collected at our site (Cabalo et al., 2025), we investigate the sensitivity of the identified gradient-ISC correlations to variations in site, scanner and stimulus. We also examine whether gradient-ISC correlations are being primarily driven by activity within certain networks. We expect a robust, reproducible relationship between these measures across movies, with differential effects in some functional networks and greater clustering at gradient poles in others. ISCs and gradient scores both enable the delineation of default mode regions from task-positive brain regions (Buckner et al., 2008; Hasson et al., 2004; Margulies et al., 2016), and in doing so, they allow us to parse out important patterns from complex spatial dynamics and inter-individual variability. By using these methods to enhance our understanding of the interactions between intersubject and inter-regional similarity, we can better define the principles that govern structure-function relationships in the brain.

## 2. METHODS

### 2.1 Gradient-ISC correlations

#### 2.1.1 Data

The main analyses were based on the Human Connectome Project (HCP) 7T data release, which includes movie-watching fMRI (Van Essen et al., 2013). From the complete dataset, which was comprised of 184 healthy adult participants (122 females, mean age 29.4 ± 3.3), 95 participants (58 females, mean age 29.5 ± 3.3) from 64 unique families were selected based on 1) whether they had complete functional and behavioral data, 2) passed motion mitigation (see Preprocessing section below), and 3) had at least 480 volumes (8 min) of functional data per run remaining after motion correction. Scans were conducted using a 7 Tesla Siemens Magnetom scanner with a Nova32 head coil at the Center for Magnetic Resonance Research at the University of Minnesota. Movie-watching data were collected during the first and fourth scanning sessions, across four sessions conducted over two days. Each scan started with a 15-min resting state sequence followed by two movie sequences, each between 15:01 and 15:21 (min:sec; TE=22.2ms, TR=1000ms; Van Essen et al., 2013). These movie sequences were composed of 4-5 different clips which were extracted from either independent films licensed under Creative Commons (e.g., a tourism film made about a small town, a mini documentary about a public garden), or Hollywood films (e.g., *Ocean’s Eleven, Erin Brokovich, The Social Network)*. Each clip ranged from 1:04 - 4:19 minutes, and was separated by 20 sec of rest. All clips started with 20 sec of rest, ended with the same 1 minute and 24 second video, and were followed by 20 sec of rest.

#### 2.1.2 Preprocessing

fMRI data underwent HCP’s minimal preprocessing pipeline (Glasser et al., 2013). This included removal of structured artifacts using independent component analysis and FMRIB’s ICA-based X-noiseifier (ICA + FIX). No slice timing correction was performed. Data was in the form of a time series of grayordinates in CIFTI format. For magnetization of the scanner to reach steady state, the first 10 volumes of each scan were discarded. Beyond the minimal preprocessing done by HCP, further steps were taken to control for potential head motion confounds. Any participant with a mean framewise displacement >0.2mm was excluded.

#### 2.1.3 Functional connectivity

After time series preprocessing, data were averaged within 1000 parcels according to the Schaefer parcellation (Schaefer et al., 2018). Parcels 533 and 903 were removed due to their small size. Time series data was concatenated across all four movie runs, and rest segments were removed to yield a total of one hour of movie-watching data per participant. Next, the mean time series for each parcel was correlated with the mean time course of all other voxels in the brain, resulting in a 998x998 symmetric correlation matrix for each subject. To create a group-averaged FC matrix, each individual matrix was z-scored using Fisher’s transformation so that the mean could be computed. This group-average matrix was subsequently converted back into r-scores.

#### 2.1.4 Gradient analysis

Individual- and group-level gradients were derived using BrainSpace (http://brainspace.readthedocs.io) *version 0.1.20*, in MATLAB *R2023b* (Vos de Wael et al., 2020). To control for noise, the input FC matrix was thresholded such that only the top 10% of correlations remain. A cosine similarity function was applied to the sparse thresholded matrix, yielding affinity matrices which describe the similarity of connectivity patterns between brain regions. Diffusion mapping was then applied to these affinity matrices to extract latent patterns from the high dimensional inputs. This method was selected over other dimensionality reduction techniques in accordance with previous works from the field (Bethlehem et al., 2020; Margulies et al., 2016; Paquola, Bethlehem, et al., 2019; Samara et al., 2023; Vázquez-Rodríguez et al., 2019). This involves normalizing the rows of the input matrix according to the diffusion operator and then solving for the eigenvalues and eigenvectors of the normalized matrix. Diffusion parameter *α* = 0.5 was used to ensure that the global properties of the dataset were maintained throughout the reduction process. This produces a low-dimensional manifold where the Euclidean distance between points represents the diffusion distance along the high-dimensional structure underlying the data (De La Porte et al., 2008). The order of the identified gradients is based on the amount of variance they explain in the original dataset.

The gradient template was generated by applying diffusion mapping to the group-averaged functional connectivity matrix. To minimize the influence of extreme values, parcel-wise gradient scores > 3 standard deviations from a subject’s median gradient score across all parcels were set to zero. Only one such outlier was identified and removed across all 95 participants. Individual gradients were computed from each participant’s functional connectivity matrix and aligned to the template using Procrustes alignment to preserve the characteristics of the individual gradients (Vos de Wael et al., 2020). Group-level gradients were then derived by averaging individual gradients scores at each parcel across subjects. The same template, generated from the discovery data set of 45 participants, was used to align both discovery and replication gradients (see Replication section) to be able to compare the results from both sets of analyses.

#### 2.1.5 Intersubject correlations

Preprocessed time series were once again leveraged to compute intersubject correlation coefficients. First, the group-averaged time series was computed by summing responses across all subjects at each timepoint and dividing by the total number of subjects. Each participant’s time series at each parcel was subsequently compared to the group mean at each parcel using Spearman’s linear correlation coefficient. This resulted in one correlation coefficient per parcel per subject. Next, the group mean ISC is computed by using Fisher’s transformation to z-score the correlation matrices, averaging the correlations for each parcel across subjects, and transforming the resulting z-scores back into r-scores. This provides one value per brain region which describes how consistently that region responded to the same movie stimulus across subjects.

ISCs can also be computed using pairwise and leave-one-out approaches. However, while pairwise methods tend to yield slightly lower correlations, we have opted here for a group mean approach due to the decreased computational intensiveness and our relatively large sample size (Nastase et al., 2019).

Once both gradient and ISC brain maps were generated, we computed the Spearman’s correlation between brain maps to determine the strength of the spatial correlation. Spearman correlation was selected here as it is non-parametric and therefore more robust with respect to the skewed distribution of ISC scores (Myers & Sirois, 2006).

#### 2.1.6 Permutation testing

Due to spatial autocorrelation and smoothing which is inherent to brain maps, we used a spatial autocorrelation-preserving spin test to assess the significance of the correlation between the ISC maps and the scores along the top three gradients (Alexander-Bloch et al., 2018). For each pair of brains maps compared (G1 vs. ISC, G2 vs. ISC, G3 vs. ISC) a Spearman correlation was performed to evaluate the spatial relationship existing between them. Then, 10,000 null correlations were generated by inflating and randomly rotating the cortical surface to yield random maps which preserve the brain’s existing patterns of spatial autocorrelation. Spearman correlations computed between spatial nulls to generate a null distribution, which was compared to the true correlations in order to assess its significance.

#### 2.1.7 Replication

To ensure that our results are both robust and generalizable, analyses were piloted in a discovery dataset consisting of 47 participants before being replicated using the same analyses methods in the remaining 48 participants. Discovery and replication groups were matched for age and sex, and related participants were kept within one of the two groups to prevent inflated similarity in the discovery and replication results due to relatedness. Results that were found to be significant in the initial analyses were compared to ensure that this significance was replicated in a new set of subjects.

### 2.2 Robustness of correlations

#### 2.2.1 Datasets

To examine the robustness of the observed spatial correlations to different datasets and movie content, we used data from two different datasets. All initial analyses were performed in the HCP 7T WU-Minn dataset (Van Essen et al., 2013). To supplement this, we also used data from our multimodal Precision Neuroimaging and Connectomics (PNI) 7T dataset (Cabalo et al., 2025), collected at the McConnell Brain Imaging Centre at the Montreal Neurological Institute. All PNI data were processed using the open-source software *micapipe v.0.2.3* (http://micapipe.readthedocs.io) (Cruces et al., 2022), and preprocessing procedures were set to approximate the HCP minimally preprocessed pipeline. This dataset currently contains movie-watching data from 13 neurotypical controls watching two three-minute clips from crime procedurals and two three-minute clips from nature documentaries for a total of 12 minutes of functional data. All participants provided free, prior and informed consent to participate. For analyses of the PNI data set, a functional connectivity matrix was generated for each subject and each movie before being averaged across movies within subjects to attempt to control for differences in scan length and number of participants in the PNI dataset.

#### 2.2.2 Robustness to stimulus content

To investigate the stability of the gradient-ISC correlations across different movies, we broke down the HCP movie-watching data into smaller clips and re-ran the main analyses. We selected two Hollywood clips, and two Creative Commons clips approximately matched for length (*Inception:* 3:48, *Erin Brockovich:* 3:51 and *Northwest Passage:* 2:23, *Welcome to Bridgeville,* 3:41). We additionally selected the test-retest clip used by HCP across each of the four movie runs. This clip is a sequence of shorter clips and had a length of 1 min and 23 sec and was played at the end of the 25-min movie run at every scan, allowing us to examine any changes in gradient-ISC relationships across repeated scans.

Functional gradients and ISCs were computed as described above. Average gradient maps were generated from the complete time series, while intersubject correlation maps were generated from each individual movie clip. Once again, Spearman correlation was used to quantify the linear relationship between gradient scores generated from all clips and intersubject correlations generated from each individual movie clip.

#### 2.2.3 Robustness to dataset

To investigate the stability of the observed correlations across datasets, we recreated the analysis completed in Aim 1, but this time correlations were computed using gradient scores and ISC maps from different datasets. Specifically, concatenated timeseries from the HCP dataset and the PNI datasets were used independently to generate average maps for both measures. These maps were then used to compute the Spearman’s correlation coefficients for all PNI gradients with the HCP ISC map, as well as for all HCP gradients with the PNI ISC map. It is important to note that these maps were generated using different movie stimuli with different numbers of timepoints, but since both measures collapse across the timeseries, this comparison can still be made and would provide strong support for the robustness of these findings.

### 2.3 Network-level correlations

#### 2.3.1 Functional networks

While previous analyses were carried out at the whole-brain level, further study was necessary to determine if macroscale correlations are being driven by brain organization occurring at the level of functional networks. To do this, we selected the 17-network functional parcellation put forth by Kong and colleagues (Kong et al., 2021). The latter has a high degree of functional granularity and allows for specific consideration of both auditory and language networks, both of which are highly relevant for the analysis of movie-watching data.

#### 2.3.2 Network correlations

After selecting an appropriate functional network parcellation, Spearman’s correlation coefficients were computed within each of the 17 networks. That is, for all networks, correlations were computed between vectors representing the gradient and ISC scores for each Schaefer-1000 parcel within that network. Only those parcels with p < 0.05 were considered. For regions that had highly significant within-network correlations across all three gradients, a mask representing the given network was put into the Neurosynth meta-analytic decoder (Yarkoni et al., 2011) to gain a better understanding of the role of that region during movie-watching.

## 3 RESULTS

### 3.1 Functional movie gradients

Diffusion map embedding, a nonlinear dimensionality reduction technique, identified the main axes of variance in FC patterns within the brain during movie-watching. In contrast with the canonical sensory-transmodal axis identified using resting-state fMRI data (Margulies et al., 2016), we identified a primary gradient (G1) anchored on one end by the somatomotor network and on the other end by transmodal systems such as default mode, frontoparietal control, and limbic networks (Figure 1B). This is consistent with previous research characterizing movie gradients (Samara et al., 2023) in that we observe maximally different gradient scores in regions responsible for somatosensory processing compared to regions associated with higher level cognitive function. The second gradient (G2) is delineated by the visual network at one end and heteromodal networks at the other, with highest scores occurring in occipital regions. Trends along the third movie gradient (G3) are generally more complex, but it has been suggested that this axis is identifying a hierarchy spanning from auditory and language-related brain regions to more heteromodal regions (Samara et al., 2023). This is also supported by distribution of G3 scores across the cortex, where we can see peaks along the superior temporal sulcus and gyrus (Figure 1B).

**Figure 1.**
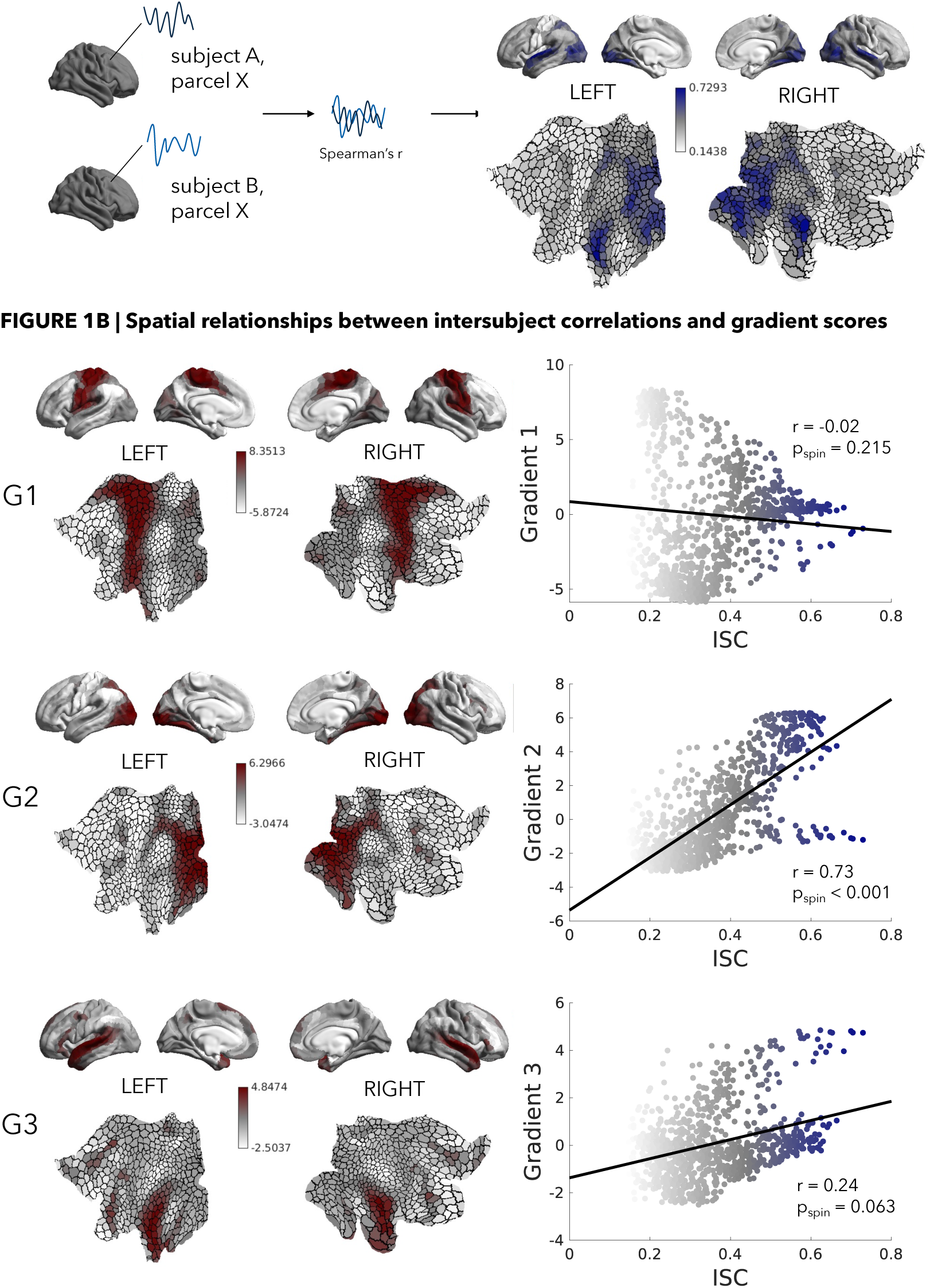
(A) Schematic representing generation of intersubject correlations (left) and their distribution on the cortical surface and flat maps (right). (B) Left column shows scores of the top three gradients mapped onto the cortical surfaces and flat maps. Right column shows correlations between gradient scores and intersubject correlations. Points are colour-coded by their intersubject correlation scores where blue represents high intersubject correlations and grey represents low intersubject correlations.

### 3.2 Intersubject correlations

Intersubject correlation analysis of fMRI time series were used to quantify the response synchronization across subjects at each parcel during movie-watching. In agreement with previous work (Hasson et al., 2004; Jääskeläinen et al., 2008; Kauppi et al., 2010; Nastase et al., 2019), peak correlations occurred in parcels located in occipital regions and along the superior temporal sulcus and gyrus, regions that are robustly stimulated by the visual and auditory features of movies (Rehman & Al Khalili, 2025; Yi et al., 2019). Frontal and primary motor regions appeared to have among the lowest ISCs, as responses in these networks are not as directly tied to the input, making them more variable across individuals (Figure 1A).

### 3.3 Relating gradient scores and intersubject correlations

To assess the spatial relationship between intersubject correlations and gradient maps, Spearman’s correlations were computed between each pair of brain maps (G1 vs. ISC, G2 vs. ISC, G3 vs. ISC). Significance of these relationships was assessed using spin permutation tests that control for spatial autocorrelation (Alexander-Bloch et al., 2018), with 10,000 permutations generated per comparison. We detected a strong linear relationship between the visually mediated G2 scores and ISC scores (*r* = 0.7, *p*_spin_ < 0.001). We see a general trend in which higher gradient scores appear to be associated with high intersubject correlations (Figure 1B). Qualitatively, this relationship appears to be driven by parcels falling within the visual network, a system that is being stimulated consistently across subjects and therefore should have highly similar fMRI responses and FC profiles. Globally, parcels within networks tend to be clustered together within a small range of ISC and gradient scores. Parcels with the highest intersubject correlations belong to the auditory network.

Relationships were weaker between ISC scores and the somatomotor-mediated G1 (*r* = 0.05, *p_spin_* = 0.215). Given that this gradient is anchored at its ends by somatomotor and default networks, both of which are not as directly and consistently stimulated during movie watching, we did not expect to see strong correlations of ISC scores with this gradient. The relationship with the auditory/language-related G3 was slightly stronger, with its moderate strength likely driven by the involvement of the auditory network during movie-watching (*r* = 0.25, *p_spin_* < 0.001).

### 3.4 Replication

Given that the G2-ISC correlation did pass the threshold for significance, these analyses were carried out again in the remaining 48 HCP participants. These participants had been held out to be used as a replication set to test the robustness of the findings. Results demonstrate a highly similar Spearman correlation between the two maps (*r* = 0.75, *p_spin_* < 0.001) with comparable significance results. The ability to replicate this finding provides support for the idea that a strong spatial relationship exists between these two maps.

### 3.5 Variability and reliability

Variability of the main gradient-ISC correlations was assessed by looking at correlations across viewings of shorter segments of the HCP dataset. This was done to evaluate whether the specific correlations observed in the main analysis were impacted by the visual or auditory content of the stimulus. Note that the gradients used here are from the entire one-hour of data collected by HCP, as gradient analyses tend to be sensitive to data amount (Samara et al., 2023), whereas the ISCs are from shorter segments. Spearman’s correlations were computed between scores from the top three gradients and ISCs for 4 short clips from the HCP dataset (*Inception*, *Erin Brockovich*, *Welcome to Bridgeville*, and *Northwest Passage*). We can see that the strong G2-ISC correlation is replicated for each individual clip, regardless of stimulus content (Figure 2). We also observe that the trend of G3-ISC correlations detected in the complete dataset is again present at the level of shorter clips.

**Figure 2.**
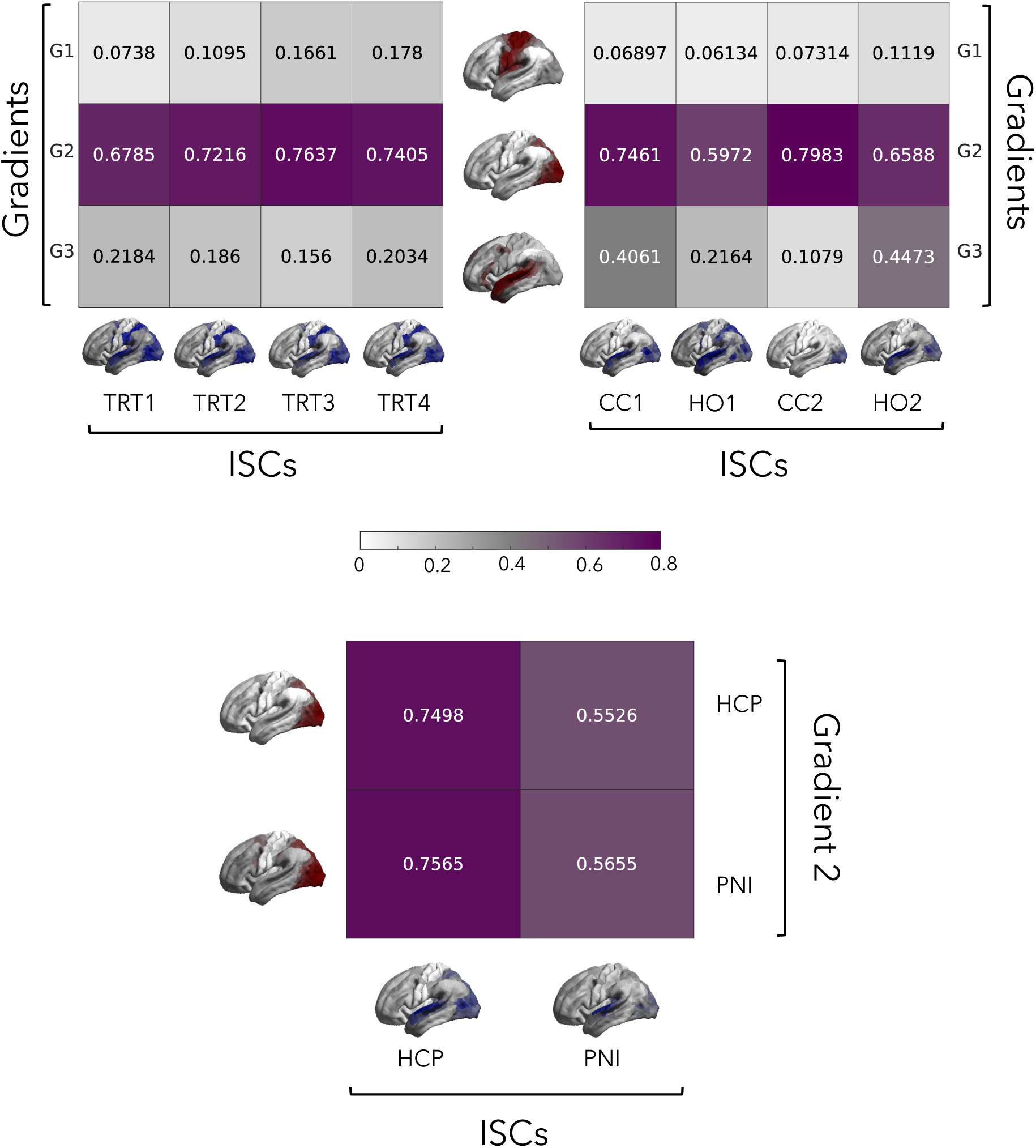
Matrices represent the Spearman’s r-values of the correlations between G2 scores and intersubject correlations across repeated viewing of the same clip (top-left) and four different clips (top-right). Darker colours represent larger correlation coefficients. Cortical maps represent the ISCs for each respective clip, as well as the gradient maps generated from the complete time series. TRT = test-retest clip, CC1 = *Welcome to Bridgeville*, HO1 = *Inception*, CC2 = *Northwest Passage*, HO2 = *Erin Brockovich*. The bottom matrix represents the Spearman’s correlations given by correlating G2 from the PNI dataset with ISCs from the HCP dataset and vice versa.

To assess reliability, the same correlations were computed, but across repeated viewing of a brief clip that was repeated during each of the four separate movie runs. The inclusion of this replication clip allowed us to assess how much this relationship varied within individuals across sessions. The correlation between gradient scores and ISCs was computed while participants watched the same 1.5-minute test-retest clip at the end of each movie run. Like the variability results, the robust G2-ISC results were replicated across each viewing of the test-retest clip. G2-ISC correlations tended to be slightly more similar across repeated viewing of the same clip compared to when watching four different clips (Figure 2). The fact that we were still able to identify this trend with such a small data amount (84 seconds for the ISC) is encouraging with respect to the reproducibility of this finding.

### 3.6 Stability

To assess the stability of these results across datasets, gradients and ISC scores from the HCP dataset were compared with their counterparts from the PNI dataset. In agreement with previous analyses, the cross-dataset gradient-ISC correlations were strongest for the second gradient. Notably, within the G2 correlations, the highest coefficients are those between G2 scores from both datasets and ISC scores from the HCP dataset (Figure 2). This suggests that the intersubject correlations generated from the HCP dataset may be more robust. This is perhaps due to the fact that the HCP dataset had a greater amount of movie-watching data per participant, or due to the slightly less engaging nature of the movies used in the PNI dataset (nature documentaries), which could decrease the robustness of the cortical synchronization between participants. Different movies constrain the sensory processing systems to different degrees, leading to fluctuations in engagement across participants and stimuli (Meer et al., 2020). Despite these fluctuations, this overall gradient-ISC relationship appears to be robust and reproducible.

### 3.7 Within-network correlations

To identify if whole-brain gradient-ISC correlations were driven by activity in particular functional networks, gradient-ISC correlations were performed within masks of the Kong 17-network parcellation (Kong et al., 2021). A few networks had particularly high correlations with ISC across the top three gradients. In particular, the Dorsal Attention B (danB) network had the highest within-network correlation for the first two gradients, and the third highest for G3. Other networks that had considerable correlations included Visual B and Default C. The presence of transmodal and default regions among those driving these whole-brain correlations was unexpected, leading us to explore the role that each area was playing during naturalistic viewing. Note that since only correlations with G2 reached the threshold for significance, only those subsequent analyses pertaining to G2 are visualized here. As shown in the right panel of Figure 3A, the R-values of these within-network correlations were finally projected back onto the cortex, and these maps were subsequently decoded by Neurosynth (Yarkoni et al., 2011) to yield key terms like ‘eye movements’, ‘occipital’, and ‘spatial information’, further supporting the idea that these correlations are being driven largely by regions involving the processing of visual and spatial information.

**Figure 3.**
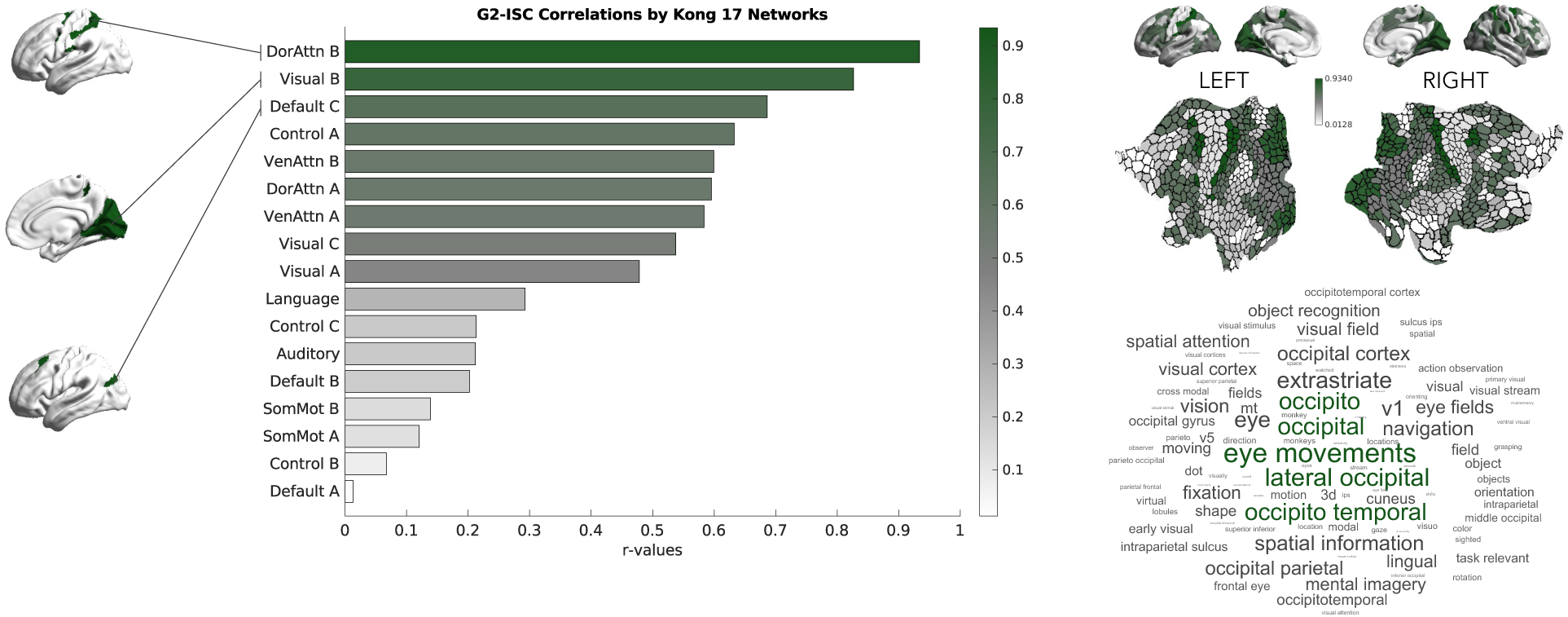
(A) bar graph represents the Spearman’s r-values of the correlations between G2 scores and intersubject correlation within each of the Kong 17 networks. Cortical surface and flatmaps were generated by projecting the within-networks r-scores back onto the cortex. This map was decoded using Neurosynth to generate the associate word cloud.

## 4. DISCUSSION

This study explored the spatial relationships between functional connectivity gradients and intersubject correlations (ISCs) under naturalistic conditions. Most fMRI research today has made use of resting state or task-based paradigms to tap into different cognitive processes. In contrast, we leveraged movie-watching as an alternative acquisition state which presents key advantages including potential decreases in-scanner head motion and enhanced identification of important individual differences by synchronizing low-level brain activity (Eickhoff et al., 2020; Finn & Bandettini, 2021; Vanderwal et al., 2017). Perhaps most interestingly, movie-watching drives the brain in a way that is more similar to what we experience in everyday life, allowing us to observe brain function under the sensory and cognitive demands of approximated ‘real world’ situations (Nastase et al., 2020; Sonkusare et al., 2019). While these naturalistic methods are gradually gaining in popularity, there is much insight still to be gleaned by using movies to capture both the controlled nature of task-based fMRI and the unconstrained nature of resting-state fMRI. As reported in previous studies, we observed increased granularity in movie gradients such that the three main axes of the naturalistic functional hierarchy are anchored by the somatomotor, visual, and auditory networks respectively (Samara et al., 2023). This granularity in gradient organization yielded by movie-fMRI provides a unique tool that can be used to probe brain organization specifically with respect to sensory systems, making them especially well-suited for the investigation of pathologies with sensory underpinnings.

Using time courses extracted from movie-fMRI, we computed ISC scores from HCP and PNI datasets, which provided 1-hour or 12 minutes and 20 seconds of data per participant, respectively. Consistent with prior work (Hasson et al., 2004; Jääskeläinen et al., 2008; Kauppi et al., 2010; Nastase et al., 2019), robust ISCs were observed in visual and auditory cortices, although overall values were somewhat lower than expected. This decrease could reflect the longer duration of HCP movies and/or the concatenation of time series data across different movie types, (*i.e.,* Hollywood and independent clips), as different movie types have been shown to evoke different levels of ISC, especially in prefrontal and higher order regions (Hasson et al., 2008)). In this way, we can understand regions that display high ISC values as being a conservative estimate of systematic, time-locked BOLD responses to the stimuli that includes only those areas that were consistently activated across a wide range of sensory and affective inputs.

We identified a significant correlation between the visually driven gradient and ISC maps generated during movie watching. This relationship was consistent across datasets, and across movie stimuli, and appeared to be largely driven by contributions from the dorsal attention network. Correlations were not as strong for the somatomotor and auditory gradients, both of which are anchored by sensory regions not driven as robustly during movie-watching. The emergence of the visual gradient as the most highly correlated of the three primary gradients is in line with our understanding of the visual system as being reliably activated during time-locked audiovisual stimulation (Hasson et al., 2004). The strong spatial relationship between these two brain maps indicates that regions such as occipital cortex, which demonstrate significant response reliability, are maximally differentiated from transmodal systems such as the default mode network. In other words, increased time-course similarity with other individuals in a given region appears to be correlated with increased functional differentiation from one’s own higher order processing networks.

Broadly, these findings reinforce previous literature describing a functional hierarchy underlying brain organization (Bernhardt et al., 2022, 2025; Hong et al., 2020; Margulies et al., 2016; Sydnor et al., 2021). These results also suggest that when the brain is active and processing complex and more ecologically valid stimuli, there is a strong correspondence between functional connectivity patterns at a whole-brain level, and response reliability across subjects at a parcel-level. What makes these results intriguing is that this represents a strong, significant and replicable correlation between a measure of between subject similarity and a measure of between region similarity. There appears to be a relationship between regions that are most differentiated along the visually mediated G2, and regions that exhibit the most reliable activation patterns across individuals. These results add an interesting perspective to our understanding of intersubject similarity and individualization: regions whose functional connectivity patterns are most dissimilar from those of the default mode networks are also most likely to be correlated across individuals. This finding is in line with recent results in the literature which indicate that this axis spanning sensory to association cortex, or reliably to idiosyncratically activated brain regions, can also be identified using different cognitive and developmental markers (Benkarim et al., 2021; Braga & Buckner, 2017; Mueller et al., 2013). In particular, the poles of the sensory-association axis can be differentiated by gradients in plasticity and heritability such that highly idiosyncratic regions in association cortex are most likely to undergo plastic changes in response to environmental variation, while sensory cortex is more closely linked to genetic factors (Sydnor et al., 2021; Valk et al., 2020, 2022). Similarly, developmental trajectories reflect this major organizational axis in that stereotyped and heritable sensory regions develop more rapidly than association cortex, where the timeline for complex, environmentally mediated plastic changes is more protracted. Increased differentiation along this sensory-transmodal axis has also been observed in healthy developmental samples (Paquola, Bethlehem, et al., 2019; B. Park et al., 2021, 2022). Additionally, recent research has suggested that this sensory-association axis is mirrored by a gradient of structure-function coupling wherein low-level sensory regions display functional activity that appears to be more closely bound by the structural constraints of the tissue, while high-level regions display a less strict relationship between these factors (Paquola et al., 2022, 2025; Preti & Van De Ville, 2019; Vázquez-Rodríguez et al., 2019). Overall, the results of this study add to a growing body of evidence that suggests that this sensory-association axis is a functionally and structurally relevant model for understanding brain organization.

Results from our reliability and stability analyses indicate that this relationship is robust and replicable across stimuli and datasets, providing support for movie watching as a brain state that does not depend on stimulus content but is defined by an organizational hierarchy which is similar to, yet distinct from that identified in resting-state fMRI. This finding builds off existing literature in the field which suggests that brain-state dynamics had higher test-retest reliability during movie-watching as compared to resting-state data (Meer et al., 2020; Tian et al., 2021; Vanderwal et al., 2017; J. Wang et al., 2017). Also supported by this finding is the idea that the pattern of brain activation observed in movie-watching is relatively independent of the content of the movie itself. While response may vary slightly with the degree to which a movie evokes sensory and emotional engagement, overall patterns are quite stable with respect to stimulus content. The consistency of these results with respect to the reliability and stability of the gradient-ISC correlations across the experimental manipulations support the concept of movie-watching as a brain state that does not directly depend on the stimulus content. Brain connectivity and BOLD-signal responses seem to emerge instead as a result of the more naturalistic way that the brain integrates not only audio and visual content, but also affective information while processing these types of stimuli. By combining the enhanced reliability of movie-watching with its increased sensory and cognitive demands, we create a tool which helps us better understand how the brain responds to the rich and dynamic environment that we experience in daily life. Traditional fMRI paradigms are highly controlled and therefore generate findings which can be difficult to generalize to real-world situations (Cantlon, 2020; Nastase et al., 2020).

By performing these analyses at the network as well as the whole-brain level, we were able to identify functional ‘sub-hierarchies’ within the cortex, providing evidence that the gradient structure used to describe macroscale functional dynamics in the brain might also be applicable within networks (Braga & Leech, 2015). These results have the possibility to deepen our understanding of the brain’s functional organization by providing insight as to how individual networks are integrated into the whole-brain organization (Margulies et al., 2016; Y. Wang et al., 2025). The results of the network-level analyses can also be interpreted in the context of systems neuroscience and related efforts to define area-level parcellations. Research has shown that parcellations generated from individual-specific resting-state data improve behavioural predictions because the relationship between intra- and inter-subject variability differs between networks, so individually distinct patterns stand out more clearly. By tapping into the tendency of sensory regions to be both minimally variable across subjects but more highly variable within subjects (Laumann et al., 2015; Mueller et al., 2013), we are able to better define functional boundaries within the brain (Kong et al., 2021).

Given that both ISC and gradient measures have been shown to be sensitive to atypical brain development, further studies could investigate the correspondence between ISC and gradient maps in psychiatrically enriched samples to understand how this relationship might be impacted. Psychiatric and neurological conditions have been shown to relate to alterations in idiosyncrasy (Benkarim et al., 2021; Dickie et al., 2018; Hasson et al., 2009; Nunes et al., 2019) and shifts in the principal gradient in resting-state data (Bernhardt et al., 2025; Dong et al., 2020; Hong et al., 2019; S. Park et al., 2021; Xie et al., 2024). How each of these measures, as well as the interaction between them are impacted in neurodivergent populations might serve to develop our understanding of the neural bases of these disorders and potentially help in efforts to discover ecologically valid biomarkers.

In conclusion, we found that there is a robust and replicable relationship between intersubject correlations scores and the second functional gradient, anchored by the visual network. The relationship persisted across different movie stimuli and datasets, suggesting that the functional organization observed during movie-watching represents a generalized brain state and not a stimulus-dependent response. We additionally found that the G2-ISC relationship was strongly driven by parcels from the dorsal attention network, indicating that the whole brain functional hierarchy identified in previous studies may also be exemplified at the network-level in the form of functional sub-hierarchies. By describing naturalistic brain organization both at the BOLD-signal level and the functional connectivity level, we obtain a more in depth understanding of the relationship between intersubject similarity and idiosyncrasy both within and across functional connectomes, allowing us to deepen our understanding of the organizational principles governing brain function.

## Acknowledgements

MS received funding from the Canadian Institutes for Health Research (CIHR). AS received funding from UBC Friedman Award for Scholars in Health and BC Children’s Hospital Research Institute Doctoral Studentship. DGC received funding from Fonds de la Recherche du Québec – Santé (FRQ-S), Quebec BioImaging Network (QBIN), Savoy Foundation, Brain Canada, and Canada First Research Excellence Fund, awarded to McGill University for the Healthy Brains for Healthy Lives (HBHL) initiative. AN received funding the FRQ-S and the CIHR. HS received funding from the CIHR. T.V. is funded by the SickKids Foundation, Health Canada via the Canada Brain Research Fund (a partnership between the Government of Canada and Brain Canada, and the Azrieli Foundation), the Canada Fund for Innovation, the Brain, Behaviour and Development Theme at BC Children’s Hospital Research Institute, and the Djavad Mowafaghian Centre for Brain Health at the University of British Columbia. BCB received support from CIHR, SickKids Foundation, Natural Science and Engineering Research Council of Canada (NSERC), Azrieli Center for Autism Research of the Montreal Neurological Institute (ACAR), BrainCanada, FRQ-S, Helmholtz International BigBrain Analytics and Learning Laboratory (HIBALL), and the Canada Research Chairs program.

## Data and code availability

The processing pipeline for the PNI dataset is part of an open-source software available on GitHub (https://github.com/MICA-MNI/micapipe). Gradient analyses were run using the BrainSpace toolbox, available on GitHub (https://brainspace.readthedocs.io/en/latest/). Additional code will be made available upon publication.

## Author contributions

MS, AS, TV, and BCB contributed to conception and study design. MS, DGC, and AN contributed to data acquisition. AS and DGC contributed to data curation and processing. MS, DGC, and AN contributed to data acquisition. MS, AS, and HS contributed to data analysis. MS, TV, BCB contributed to drafting the manuscript and preparing the figures.

